# DIRECTED GRAPH THEORY FOR THE ANALYSIS OF BIOLOGICAL REGULATORY NETWORKS

**DOI:** 10.1101/2023.10.02.560622

**Authors:** Martha Takane, Saúl Bernal-González, Jesús Mauro-Moreno, Gustavo García-López, Bruno Méndez-Ambrosio, Francisco F. De-Miguel

**Author notes:** Corresponding author: Martha Takane Instituto de Matemáticas, Universidad Nacional Autónoma de México.

## Abstract

Regulated biological networks are commonly represented as logical diagrams, in which the exact interactions between the elements remain out of sight. Here we propose a new type of excitation-inhibition graph based on Boolean logic, which we name “logical directed graph or simply, logical digraph of the biological system”. Such logical digraph allows the representation of every possible regulatory interaction among elements, based on Boolean interactions. The logical digraph contains information about the connectivity, dynamics, limit cycles, and attractors of the network. As proof of the application, the logical digraph was applied to analyze the functioning of the well-known neural network that produces oscillatory swimming in the mollusk Tritonia. Our method permits to transit from a regulatory network to its logical digraph and vice versa. In addition, we show that the spectral properties of the so-called state matrix provide mathematical evidence about why the elements in the attractors and limit cycles contain information about the dynamics of the biological system. Open software routines are provided for the calculations of the components of the network and the attractors and limit cycles. This approach offers new possibilities to visualize and analyze regulatory networks in biology.

## INTRODUCTION

Regulatory networks in biology are established when a collection of elements, namely molecules, neurons, or individuals, interact with each other to determine function. The time-dependent interactions of the components of a regulatory network define characteristic operation cycles, which may span from fractions of a second to several days. Detailed experimental explorations of the workings of regulatory networks can be enriched by mathematical models that reproduce the behavior of the network and contribute to the design of new hypotheses and experiments. In this article, we provide a quantitative method based on Boolean algebra to reproduce behaviors and missing elements of the network.

Regulatory networks are commonly studied by representing interactions among elements in the network with an excitation-inhibition graph, see [1,2,3,4,5,6,7,8,9]. For example, *a* and *b* denote two interacting elements, each having a binary Boolean behavior. We may now suppose that activation of *a* turns *b* on. Such activation, or excitation, can be described as *a* → *b*. If by contrast, *a* inhibits *b*, the expression will be *a*⊣ *b*. Such a simple description can now be enriched in Boolean terms by using 1 to describe an active state and 0 to describe an inactive or inhibited state. However, such notation is insufficient to explain more sophisticated interactions. For example, gene *a* may be activated when gene *b* is inhibited. Alternatively, gene *a* may remain invariant upon activation of gene *b*. Such and other interactions naturally occurring in nature cannot be described solely by the excitatory and inhibitory connectives. Another method to describe networks and circuits in biology is using the logical electrical diagrams used in engineering. However, they act as black boxes in which the exact interactions between the elements remain out of sight, see [5].

To overcome such limitations, here we propose a digraph (directed graph) that we will call a ***logical digraph*** of the biological system. The logical digraph can be built using eight logical connectives with their combinations, which represent every possible interaction occurring among any two elements. When combined with appropriate Boolean functions the digraph accurately defines the dynamics of the biological regulatory system. In addition to being algorithmizable, our method reports on the system’s attractors and limit cycles, which correspond to the Perron-Frobenius eigenvectors of the ***state matrix*** that describes the transition from one state to another over time. Therefore, the state matrix contains information on the dynamics of the biological network (As a general reference we recommend [10] Stuart A. Kauffman’s “The Origins of Order. Self-Organization and Selection in Evolution”). As new concepts appear along the paper, they are applied to the simple neural circuit that controls swimming in the mollusk Tritonia to exemplify the use of the logical digraph.

### Glossary

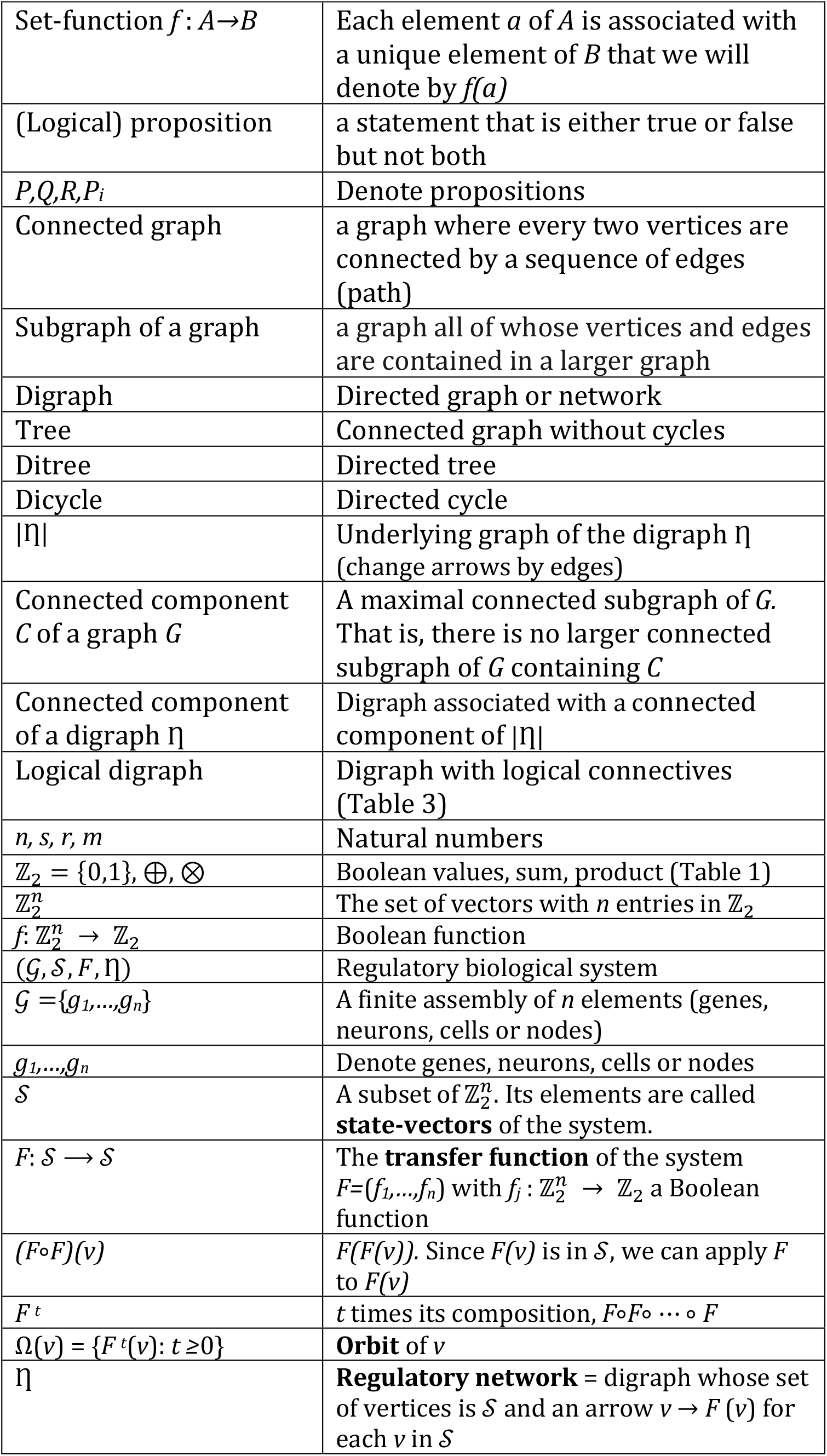
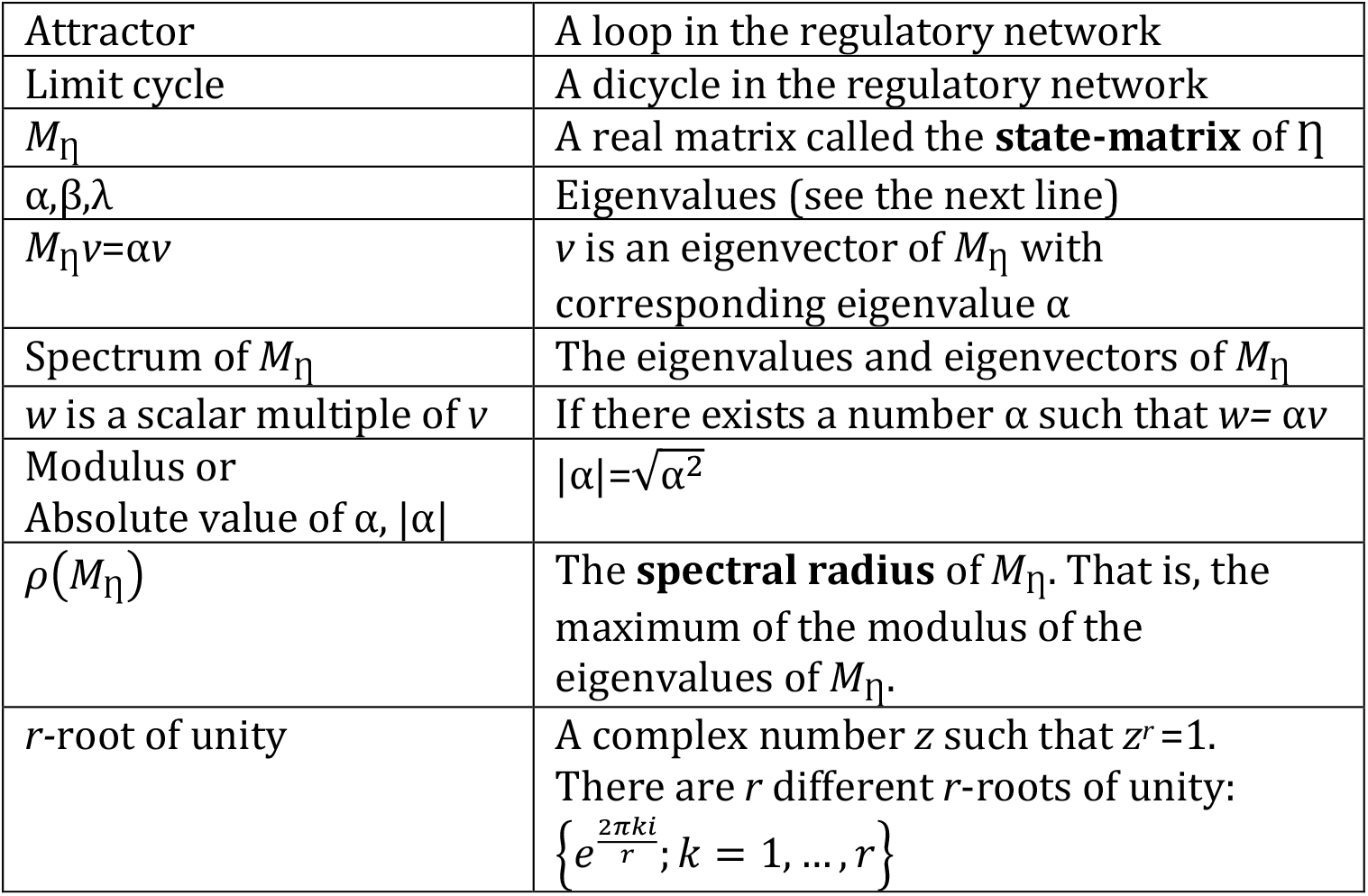

### I. THE LOGICAL DIGRAPH OF A REGULATORY BIOLOGICAL SYSTEM. DEFINITIONS AND BASIC NOTIONS

A regulatory biological system and its dynamics may be described by the quartet of symbols (𝒢, 𝒮, *F*, η). The symbol, 𝒢 ={*g*_*1*_,*…,g*_*n*_} is a finite assembly of *n* elements (genes, neurons, cells or nodes), each of which acquires either the Boolean active state (on, true) and a 1 value, or the inactive state (off, false) and a 0 value. We will denote the set of Boolean values as 𝕫_2_ ={0,1}, together with their usual Boolean operations, addition ⊕ and multiplication ⊗, described in Table 1.

**Table 1.** Boolean values of the Boolean sum and multiplication of two interacting elements in a regulatory network.

|  |  |  |
| --- | --- | --- |
| $\oplus$ | 0 | 1 |
| 0 | 0 | 1 |
| 1 | 1 | 0 |

|  |  |  |
| --- | --- | --- |
| $\otimes$ | 0 | 1 |
| 0 | 0 | 0 |
| 1 | 0 | 1 |

In the set 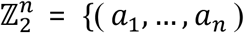: *a*_*i*_ ∈ 𝕫_2_ for each *j*=1,…,*n*} the *j*-th coordinate corresponds to the element *g*_*j*_ of 𝒢. This set contains all the possible states (0,1) of the elements of 𝒢, which we will call **state-vectors**. All the state-vectors of the biological system are contained in 𝒮, which is a subset of 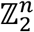.

The symbol **η** represents a digraph whose vertices are the state vectors (the elements of 𝒮) and represents the regulatory network. η contains information about the time-evolution of the regulatory cycle. In (*a*_1_, …, *a*_*n*_) → (*b*_1_, …, *b*_*n*_), the arrow means that the state (*a*_1_, …, *a*_*n*_) changes to the state (*b*_1_, …, *b*_*n*_) in a unit of time. Therefore, it may be represented as (*a*_1_, …, *a*_*n*_) → (*b*_1_, …, *b*_*n*_). These changes are given by the **transfer function** *F=*(*f*_*1*_,*…,f*_*n*_) of the system, defined by *n* Boolean functions *f*_*j*_ : 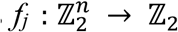 that describe how the elements of 𝒢 act on the element *g*_*j*_. In other words, the transfer function *F* goes from 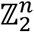 to 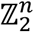 and an arrow (*a*_1_, …, *a*_*n*_) → (*b*_1_, …, *b*_*n*_) in η means a transfer function (*b*_1_, …, *b*_*n*_) = *F*(*a*_1_, …, *a*_*n*_).

### LOGICAL CONNECTIVES

We may now go further using logical connectives that express excitation and inhibition. To start, we will use again the excitation connective to explain the essentials of the use of connectives:

#### Excitation

An active *a* (*a*=1) induces the transition of *b* from inactive (*b*=0) to active (*b*=1). The binary value 1 models excitation from *a* to *b*, which is represented by an arrow *a*→*b*. Table 1 shows that the excitation value depends on the initial states of either component. If *a* is off (*a*=0), *b* will remain inactive (*b*= 0). Note that if *b* is initially active, the activation of *a* also renders a 1 value. By contrast, the initial value of *b* remains unchanged if *a* = 0.

#### Inhibition

Direct inhibition means that activity of *a* turns *b* off. By substituting the *a*=1 value to indicate that *a* is active, we have that *b* = 0. Alternatively, if *a* is inactive it does not affect *b*. Therefore, in such a situation, *a* = *b*.

Table 2 shows that the excitation and inhibition connectives expressed as a Boolean function may each produce four possible results, depending on the initial states of *a* and *b*.

**Table 2.** State-dependent Boolean values upon excitation and inhibition.

| $(a,b)$ | $a \rightarrow b$ | $a \vdash b$ |
| --- | --- | --- |
| (1,1) | 1 | 0 |
| (1,0) | 1 | 0 |
| (0,1) | 1 | 1 |
| (0,0) | 0 | 0 |

### MORE LOGICAL CONNECTIVES

The complexity of biological interactions goes far beyond excitation or inhibition. The diverse ways by which *a* may affect *b* depend on the momentary states of each interactive element. Furthermore, one must consider the global activity even when *a* is inactive, a fact not always considered in this type of analysis. For such reason, six additional connectives need to be added to accurately describe dynamics or real biological regulatory networks:

#### Negation of excitation (*a* ⤏ *b*)

A first case occurs if activation of *a* inhibits an initially active *b*, that is (1 ⤏ 1) = 0. In a second case, *b* remains inactive during the activity of *a*, (1 ⤏ 0) = 0. Both cases resemble the inhibition seen above. However, here the inactivity of *a* produces inactivation of *b*, (0 ⤏ *b*) = 0. Alternatively, the initial inactivity of both *a* and *b* produces activation of *b*, (0 ⤏ 0) = 1.

#### Negation of inhibition (*a*− ⊣ *b*)

As opposed to the previous case, any difference in the initial activity between *a* and *b* results in a change in the *b* value. However, inactivity in both *a* and *b*, keeps *b* inactive.

#### Implication (*a* ⟹ *b*)

The activity in both *a* and *b*, keeps *b* active. By contrast, the inactivity of *a* produces spontaneous activation of *b*; in the other cases, *b* will remain inactive.

#### Negation of implication (*a* > *b*)

As opposed to the implication, the only case where *b* is activated is when *a* is active and *b* is inactive.

#### Disjunction 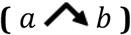

Simultaneous activity of *a* and *b* is necessary to maintain the activity of *b*.

#### Negation of disjunction 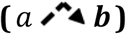

The unique case when *b* becomes inactive is in simultaneous activity of *a* and *b*.

### COMBINING LOGICAL CONNECTIVES

Up to now, the interactions between two elements have been defined by a single symbol. The following eight complex interactions will be defined by a combination of two or more logical connectives joined with the logical connective “AND” (or disjunction).

#### Doble implication or If and only if

(Means *a* ⟹ AND *a*− ⊣ *b*. That is, *b* will be active if *a* and *b* have the same value.

#### Negation of the Doble implication 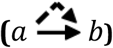

Means 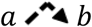 AND *a* ⟶ *b*. That is, alternating the values of *a* and *b* will activate *b*.

#### “*a*” as an action from *a* to *b* 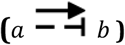

Means *a* ⟶ *b* AND *a* − ⊣ *b*. That is, regardless of its value, *b* takes the value of *a*.

#### “NOT

*a*” (as an action from *a* to *b*) or **Negation of** *a* 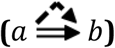 Means 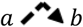 AND *a b*. That is, regardless of the value of *a, b* takes the opposite.

#### “*b*” (as an action from *a* to *b*) 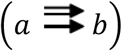

Means *a* ⇒ *b AND a* → *b*. That is, the value of *b* remains invariant.

#### “**NOT** *b*” (as an action from *a* to *b*) 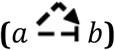

Means 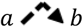 AND *a b*. That is, *b* takes the value opposite to its original value.

#### Tautology

(as an action from *a* to *b*) 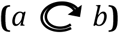, *b* always will be activated.

#### Contradiction

(as an action from *a* to *b*) 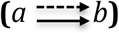. Means *a* ⤏ *b* AND *a* → *b*, meaning that *b* will always be inactivated.

Table 3 summarizes the connectives in terms of their logical propositions, boolean functions, logical propositions and values. The strategy now is that for each Boolean function in *n* variables, we will build the logical proposition it defines and the corresponding logical digraph, and vice versa.

**Table 3.**
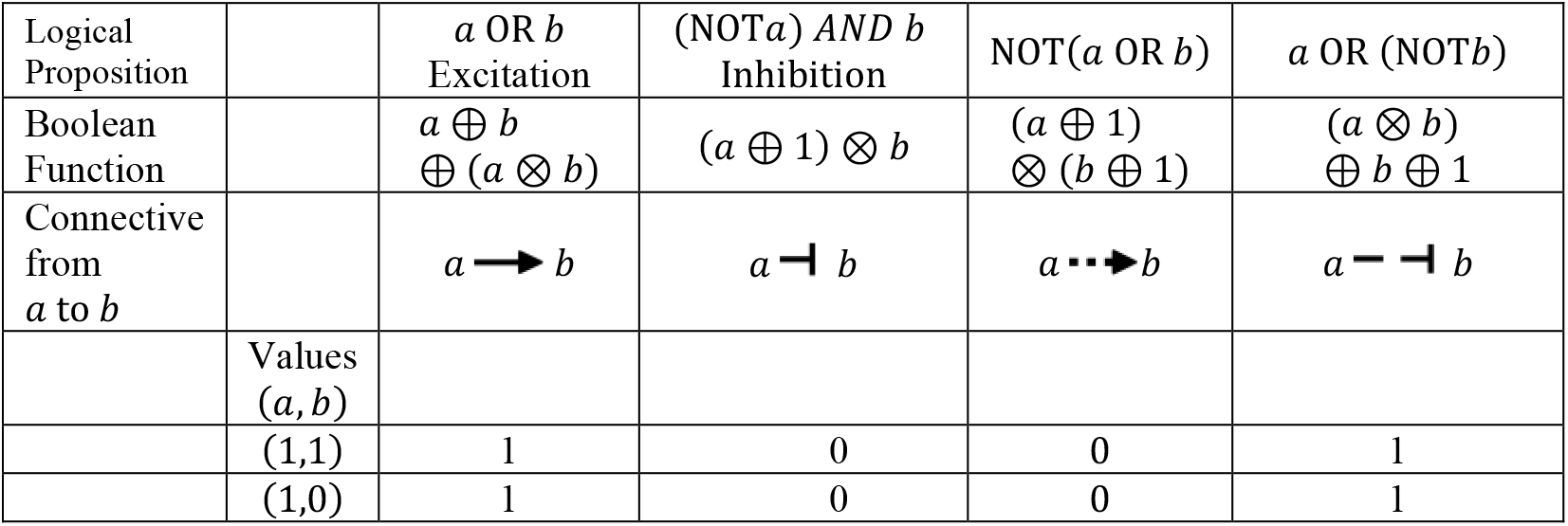
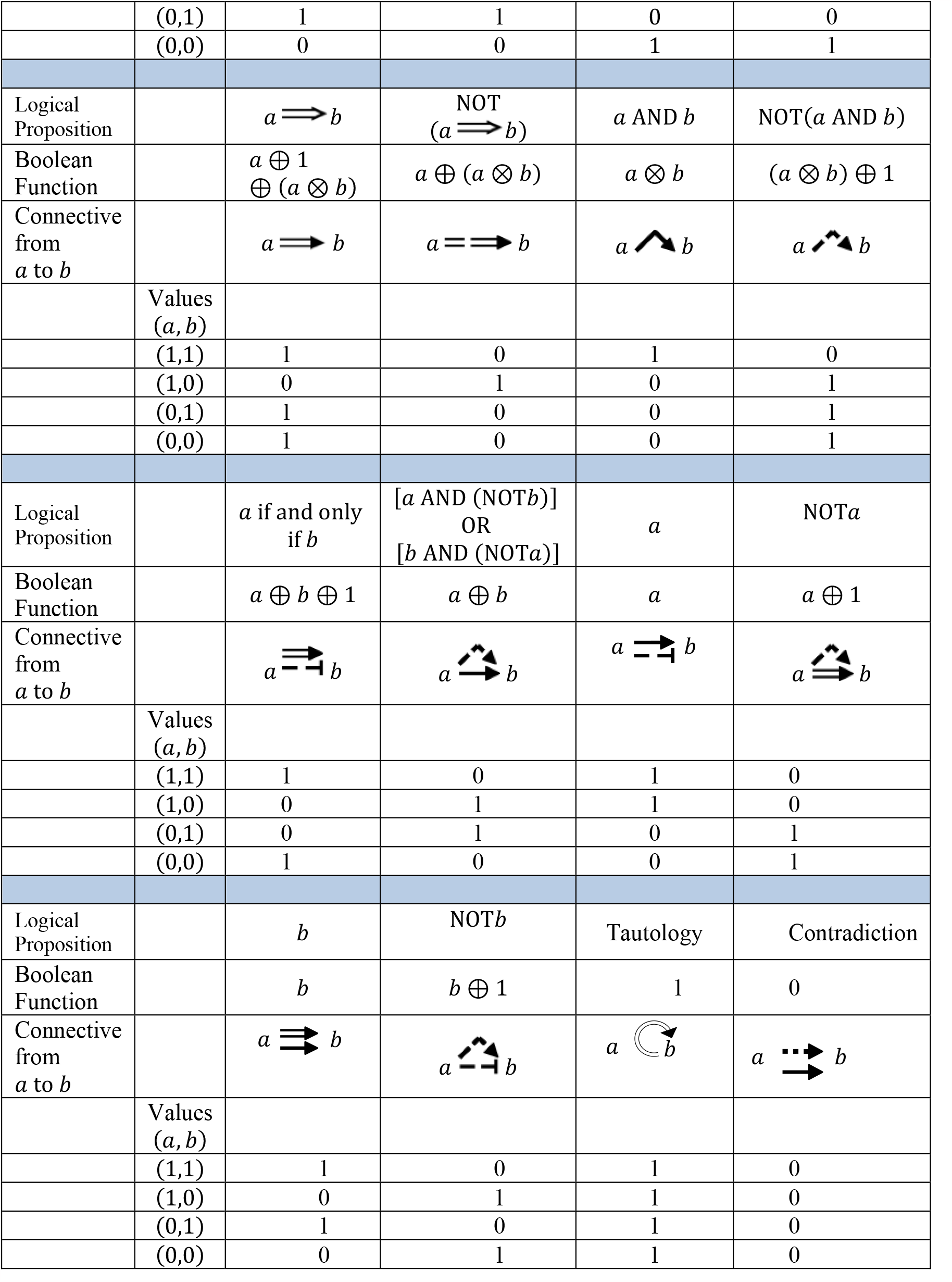
Connectives, logical propositions, Boolean functions and values of the 16 actions of *a* onto *b*.

### BIJECTION BETWEEN LOGICAL PROPOSITIONS AND BOOLEAN FUNCTIONS

To represent the regulatory network in Boolean terms we need a conversion from logical propositions to Boolean functions. Such a connection can be achieved if there is a tautology (bidirectional equivalence), meaning that the conversion is true in every possible interpretation. For example, the two logical propositions *P* and *Q* are equivalent if the logical proposition “*P* if and only if Q” is a tautology. Table 3 already introduced the bijections between symbols, logical propositions and Boolean functions. Table 4 shows that the true table of the connective excitation is the same as the logical proposition OR, and their Boolean logical propositions are also the same. Therefore, there is a tautology and we can substitute one for the other.

**Table 4.** Example of equivalences between logical and Boolean propositions.

| Values<br>$(a,b)$ | $a$ OR $b$ |
| --- | --- |
| (1,1) | 1 |
| (1,0) | 1 |
| (0,1) | 1 |
| (0,0) | 0 |

Table 5 shows a bijection between the set of logical propositions in *n* variables and the set of Boolean functions from 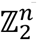 *to* 𝕫_2_, for each natural number *n≥*1.

**Table 5.**
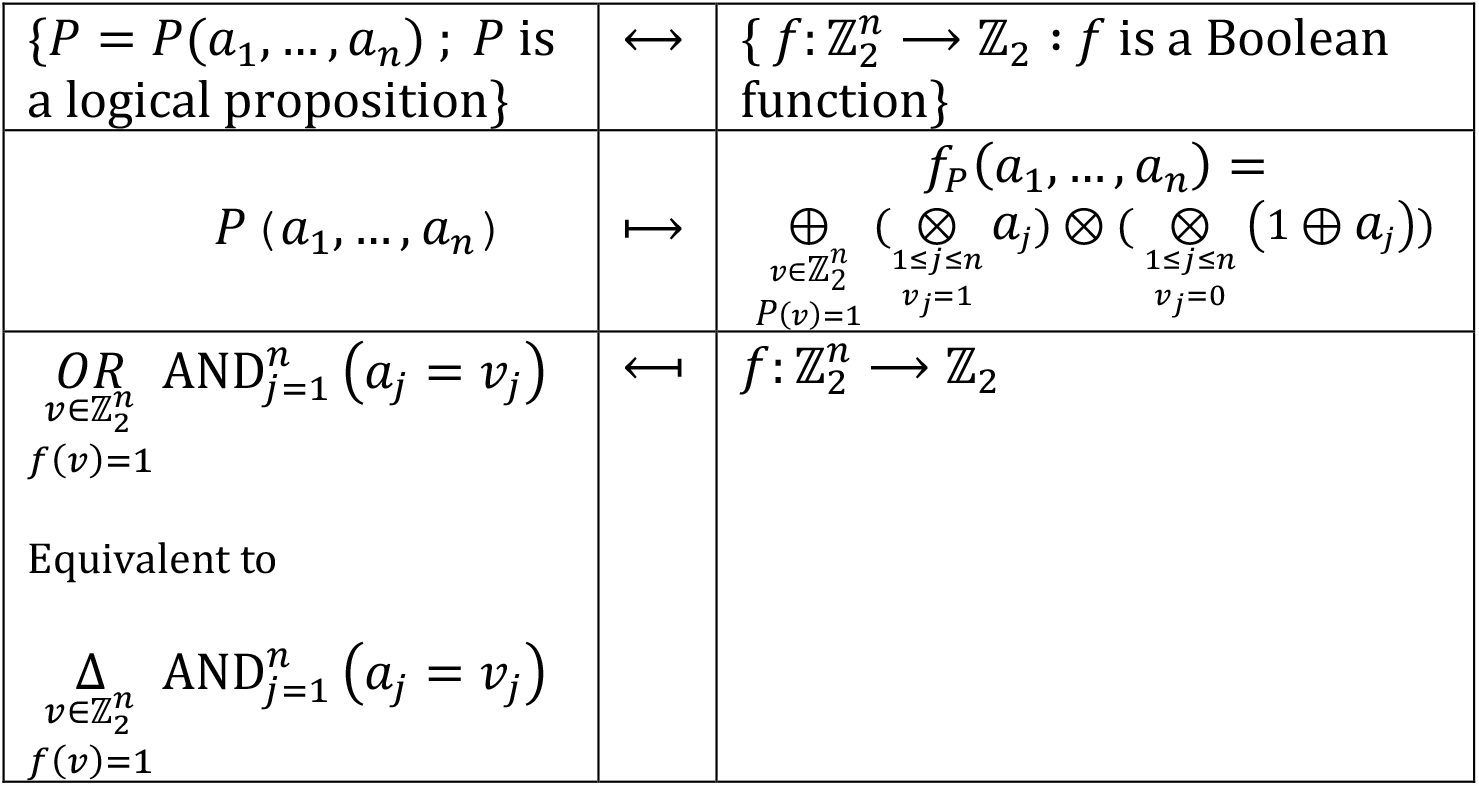
Examples of bijections between a set of logical propositions in *n* variables and the corresponding set of Boolean functions.

### FROM A REGULATORY BIOLOGICAL SYSTEM TO ITS LOGICAL DIGRAPH AND VICE VERSA

We will now return to the biological regulatory system (𝒢, 𝒮, *F*, η) and will use η to build the transfer function *F* (alternatively, we could use *F* to build η). The supplementary material contains a computer routine that calculates the function having *F* from a predefined regulatory network η.

The logical digraph of a biological regulatory system (𝒢, 𝒮, *F*, η) has one *n* vertex for each element of 𝒢. Such vertices will be denoted as *g*_1_, …, *g*_*n*_ (𝒢 = {*g*_1_, …, *g*_*n*_}), and the directed connectives joining the vertices correspond to the logical connectives in Table 3. 𝒮 is the set of state-vectors and *F* = (*f*_1_, …, *f*_*n*_) is the set of Boolean functions which define the regulatory network η.

It is now necessary to describe each Boolean function 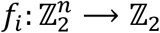 in terms of the sum and multiplication of 𝕫_2_, such that:

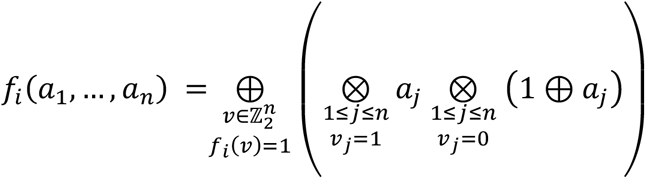

To obtain its corresponding logical proposition *P*_*i*_ we will apply the bijection to each Boolean function *f*_*i*_ of *F* . From Table 3 we will obtain the logical connectives for every element in 𝒢 (including *g*_*i*_) to the vertex *g*_*i*_ . Remember that the logic digraph is built with all the interactions among all the genes, including those established by a gene onto itself.

### APPLICATION OF THE LOGICAL DIGRAPH TO ANALYZE THE NEURAL NETWORK OF SWIMMING IN A MOLLUSK

The well-known and relatively simple neural network controlling swimming of the mollusk Tritonia offers numerous advantages for testing mathematical network theory applied to animal behavior (see [11]). As in other invertebrates, swimming in Tritonia is produced by a sequence of sigmoidal body waves upon alternate cycles of contraction of ventral and dorsal muscles. The beauty of such neuronal circuitry relies on the combination of its simplicity and the reproducibility of the emerging behavior. Studies made by Getting, Katz and others (see [1,12,13,14,11, 15]) have shown that swimming is produced by a central pattern generator integrated by four types of central neurons that establish a stereotyped connectivity.

Motoneurons carry the output information, which alternates the contraction of dorsal and ventral muscles. In brief, activation of the Dorsal Interneuron (DI), generates bursts of action potentials that activate via excitatory connection, the dorsal motoneurons. Swimming starts in the upward direction. Simultaneously the DI neuron activates the cerebral type 2 (C2) neuron. The C2 neuron responds with a stream of impulses that in turn produce a delayed excitation of the ventral interneurons (VI). Two types of VI neurons cooperate to the pattern, however, for the purpose of this study, it is sufficient to condense both as a single representative neuron (see also [16]). The VI interneuron connects to the ventral motoneurons that produce ventral contraction of the animal [17].

Up to here, only direct excitatory and inhibitory connections have been defined. However, the timing of the cyclic activation of the CPG requires additional connections. Reciprocal inhibitory connections between the DI and VI motoneurons are essential. The DI and VI interneurons communicate with each other via reciprocal inhibitory connections. The activity of DI neurons silences VI neurons, cleaning the circuit to produce a sole dorsal output, while extending the excitation time of the dorsal motoneurons. However, the excitation of C2 neurons followed by the excitation of VI neurons inhibit DI and C2 interneurons, allowing a sole ventral output of the circuit. A third type of connection contributing to the duration of the burst of impulses by the DI and VI motoneurons are excitatory autapses, namely, synaptic connections that neurons establish onto themselves. Activation of autapses during the bursts of action potentials elongates excitation of the DI or VI neurons, elongating the duration of their firing and producing inhibition of the antagonistic neuron.

From the information listed above, the dynamics of the neural network of Tritonia swimming with three neuron types firing orderly can be represented as a sequence of steps shown in Figure 2, where the contribution of neurons follows the order (ID, IV, C2).

**Figure 1.**
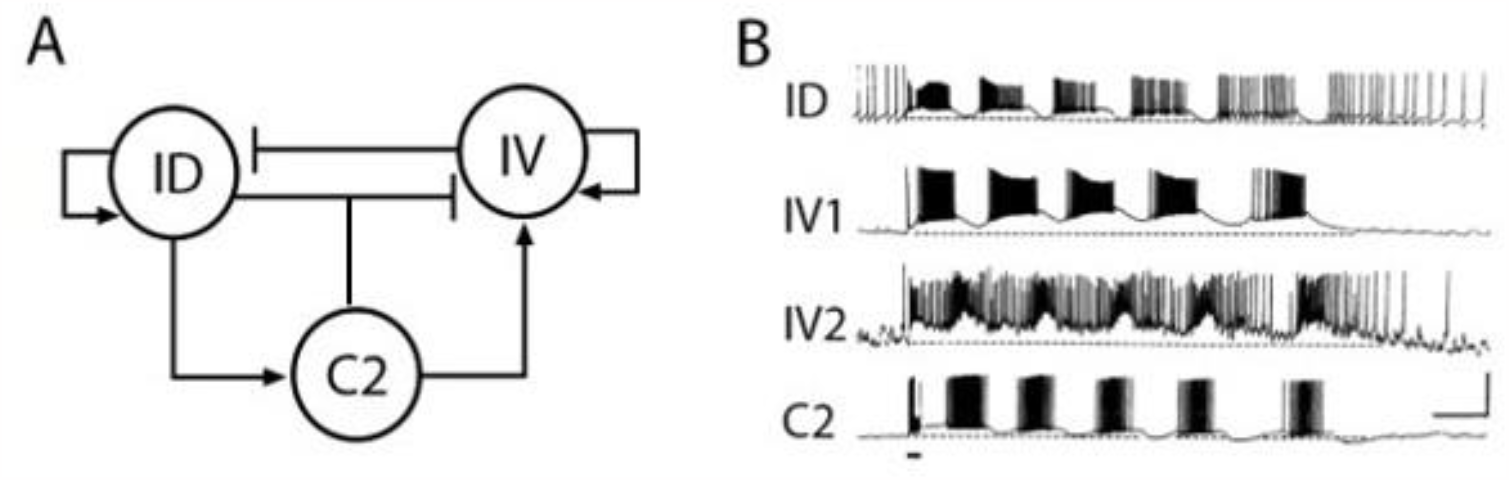
The neuronal circuit of swimming in Tritonia. (A). Basic neuronal circuit that integrates the CPG. DI, dorsal interneuron; VI, ventral interneurons; C2, cerebral neuron type 2. The DI and VI interneurons connect directly to their respective motoneurons (not shown). Excitatory connections are indicated with the vertical small bars; inhibitory connections are indicated with circles. The small bar below indicates the swimming-initiating stimulus. (B). Simultaneous intracellular recordings from each type of neuron. Note the phase differences in the firing of the different neuron types. Adapted from [16].

**Figure 2.**
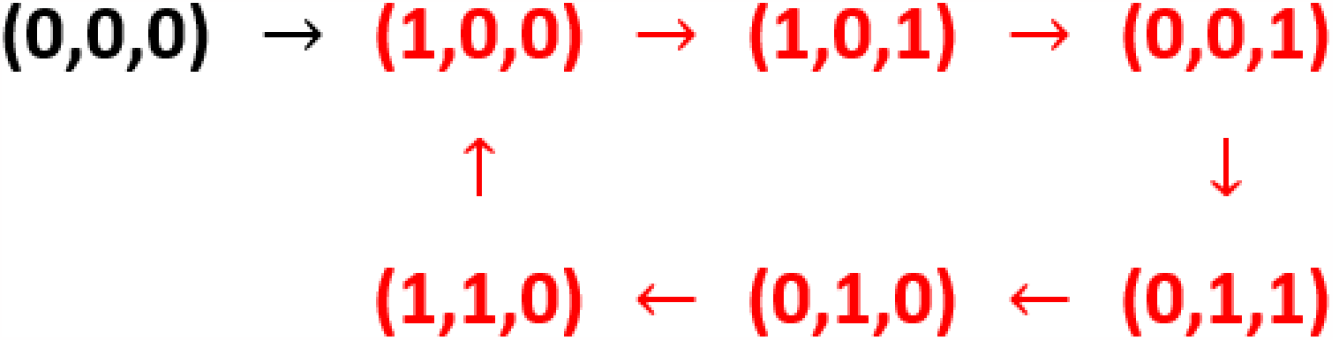
State vector representation of the sequence of neuronal activation leading to Tritonia swimming. Each parenthesis contains the state-dependent “on” (1) or “off” (0) values characterizing activity or rest of (ID, IV, C2). The arrows indicate the activation sequence, which repeats itself for several cycles once started. The dicycle of the network is in red, see (II.4).

### CONSTRUCTION OF THE TRANSFER FUNCTION *F*

We are now ready to build the Boolean function 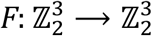, which in this case is defined as *F* = (*f*_*DI*_, *f*_*VI*_, *f*_*C*2_) and its logical digraph. We must remember (see the glossary) that the composition *F*^*t*^ gives the state of the network at time *t*, so for a large enough *t* (*t* ≥ 8, in this example), the biological system will transit around the whole dicycle. For references of Boolean functions and set theory, see [19,20,21].

We will start by defining:

1. 𝒢 ={ID, VI, C2}
2. 𝒮 = {(0,0,0), (1,0,0), (1,0,1), (0,0,1), (0,1,1), (0,1,0), (1,1,0)}, which is the set of the state-vectors that represent the different working states. Recall that η is obtained by applying the transfer function *F* to each vector of 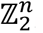, and that there is a unique arrow from *v* to *F*(*v*), for every 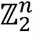. Therefore,
3. From the network η, we know that:

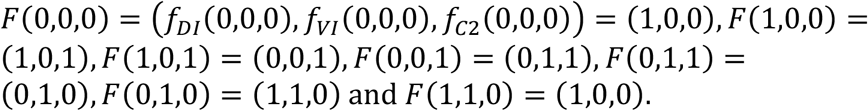

Where,

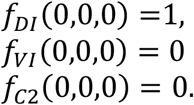

The same procedure must be repeated for every state vector of the system.

To build explicitly the Boolean function 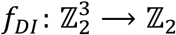 we have to find all the state-vectors of 𝒮 that under *f*_*DI*_ are going to give 1. They are (0,0,0), (1,0,0), (0,1,0) and (1,1,0).

By using bijection, we have:

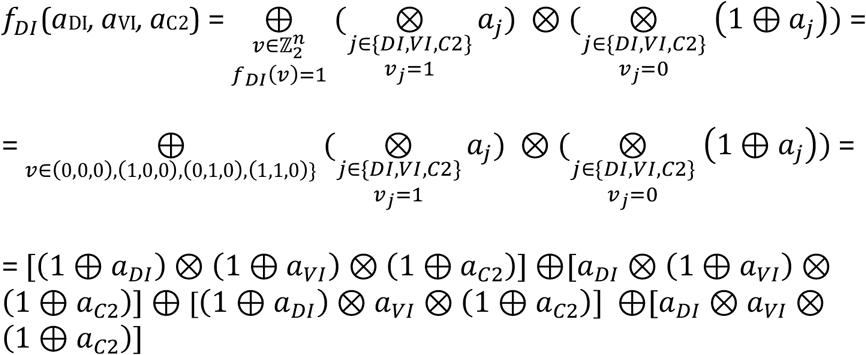

And now, by using the known equalities

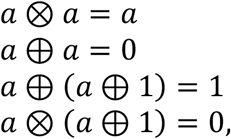

we get that *f*_*DI*_(*a*_DI_, *a*_VI_, *a*_C2_) =1⊕ *a*_C2_.

To build *f*_*VI*_, we repeat the procedure. The state vectors with a 1 value under *f*_*VI*_ are (0,0,1), (0,1,1) and (0,1,0). Then,

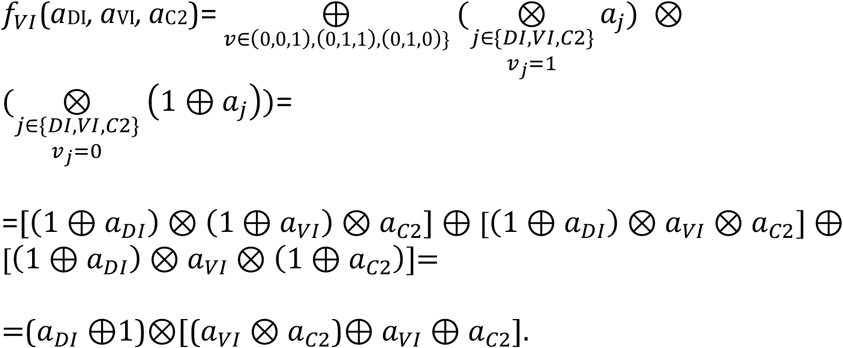

Likewise, *f*_*C*2_(*a*_DI_, *a*_VI_, *a*_C2_) = 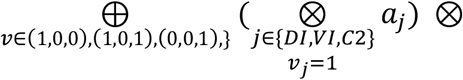

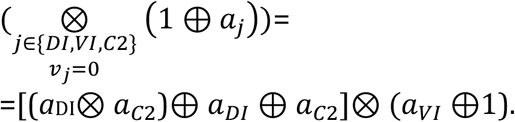

Therefore,

*F* (*a*_DI_, *a*_VI_, *a*_C2_) = (1⊕ *a*_C2_, (*a*_*DI*_ ⊕1)⊗[(*a*_*VI*_ ⊗ *a*_*C*2_)⊕ *a*_*VI*_ ⊕ *a*_*C*2_], (*a*_DI_ ⊗ *a*_*C*2_)⊕ *a*_*DI*_ ⊕ *a*_*C*2_]⊗ (*a*_*VI*_ ⊕1)).

Again, let’s take *f*_*DI*_ (*a*_DI_, *a*_VI_, *a*_C2_) =1 ⊕ *a*_C2_ to find the logical connectives in DI. Table 3 shows that the logical proposition corresponding to 1⊕ *a*_C2_ is “NOT(C2)”, which corresponds to:

C2 DI.

Let’s now repeat the analysis for VI:

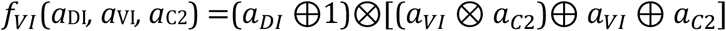

with the corresponding logical proposition “NOT(DI) **AND** (VI OR C2)”. From Table 3 we obtain the logical digraph in VI:

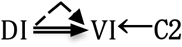

Finally, we obtain C2:

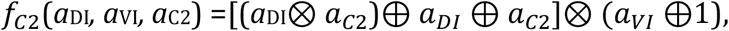

Whose logical proposition is “(DI OR C2) **AND** NOT(VI)”, and its logical digraph is:

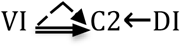

The assembly of all the pieces above provides the logical digraph of the neural regulatory network of Tritonia swimming shown in Figure 3:

**Figure 3.**
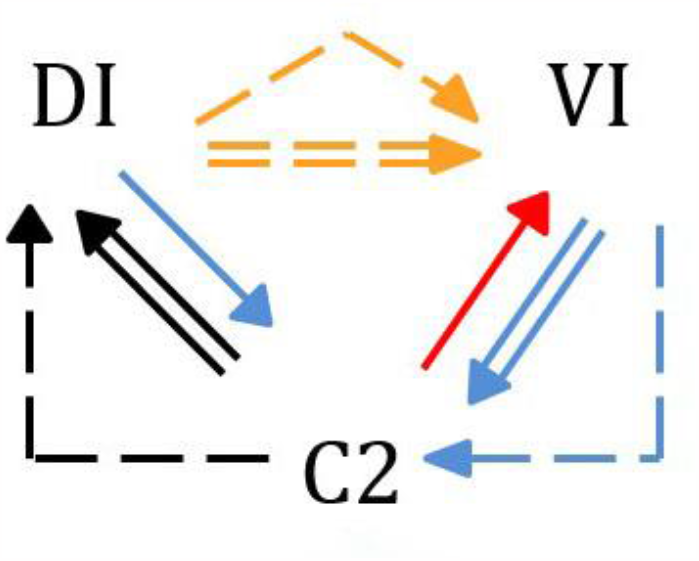
Digraph describing the neural network of Tritonia Swimming.

Figure 4 shows that if an alternative, we consider solely the excitation-inhibition graph of the biological system of Tritonia swimming (Figure 1B), the resulting network fails to reproduce the functional network.

**Figure 4.**
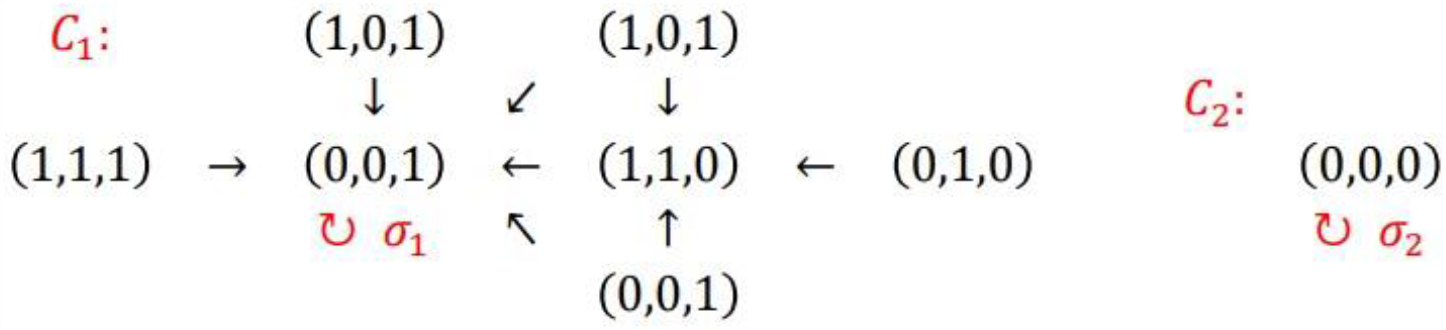
Regulatory network of Tritonia swimming based on pure excitation and inhibition connectives. The network has two connected components *C*_*1*_ and *C*_*2*_. The loop σ_1_ in (0,0,1) is the attractor of *C*_*1*_ and the loop σ_2_ in (0,0,0) is the attractor of *C*_*2*_. (see also sections II.4 and III).

### PART II. DYNAMICS OF THE NETWORK

In the first part of our paper, we went from the transfer function *F*: 𝒮 ⟶ 𝒮 of a biological system to its logical digraph. In this section, we will discuss the tools for studying the dynamics of a regulatory network of a biological system.

#### Sketch

To describe the connected components of η and the dynamics of the biological system, in this section we will use the transfer function *F*, its orbits and the state-matrix *M*_η_ of the regulatory network η.

Section **II.0** contains certain necessary definitions to understand the network topology and the flow of information between *F* and η.

Section **II.1** defines the basic properties of the transfer function *F*. Section **II.2**, defines the properties of the **state-matrix *M***_η_ and its

**spectrum**, composed of the eigenvectors and eigenvalues of *M*_η_. The Spectrum gives us additional tools to study the dynamics of the regulatory network η.

Section **II.3** shows how to translate information between the transfer function *F* and the state-matrix *M*_η_. This transfer of information will help us to retrieve the Boolean information of the network when we study the dynamics of the network over time.

Section **II.4**. A regulatory network η is a digraph, that is, a graph that instead of edges has arrows, see Figures 2, 4 and the glossary for definitions. η may be connected, as in the network in Figure 2, or not connected as in Figure 4, where η has two connected components *C*_*1*_ and *C*_*2*_. It is important to know if the network η is connected or not, because each connected component contains different biological information. We will have to build and study each connected component separately to avoid information mixtures that will render false results.

The supplementary material **SM1** contains computer programs designed to determine which components are connected in η; The supplementary material **SM2** contains a computer program designed to obtain the transfer function *F* from the regulatory network η.

#### II.0. Topology of network dynamics

Let us begin this second part by introducing some necessary definitions to analyze the network dynamics. Figure 6 shows the essential configurations of the network. A simple case occurs when one element connects to itself forming a loop, which is called an **attractor** (Figure 6a). Mathematically, an **attractor** *v* is a fixed point of *F*. Therefore, *F*(*v*)=*v*. The network in Figure 4 has two attractors, one at the state vector (0,0,1) and another at the state vector (0,0,0).

**Figure 5.**
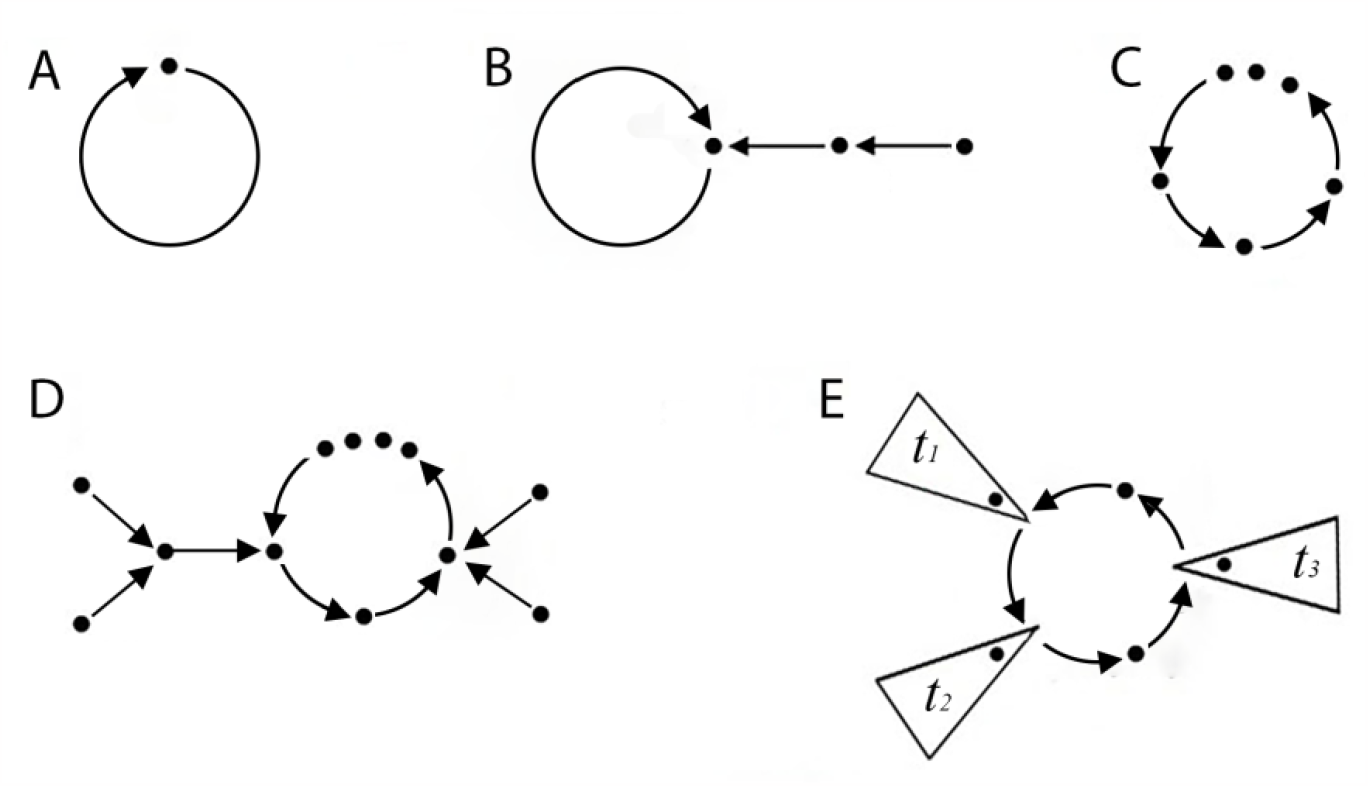
Elementary shapes of a connected network η. Each paragraph describes a connected network (digraph). The dots are elements and the arrows are connectives (A) Single connected element a loop (attractor) without ditrees. (B) An attractor with a ditree. (C) Structure of a dicycle (limit cycle). (D) A limit cycle receiving two ditrees (on the right and on left side). (E) Simplified representation of a limit cycle with ditrees represented as *t*_1_,…, *t*_*l*_. Each ditree may contain one or multiple elements.

**Figure 6.**
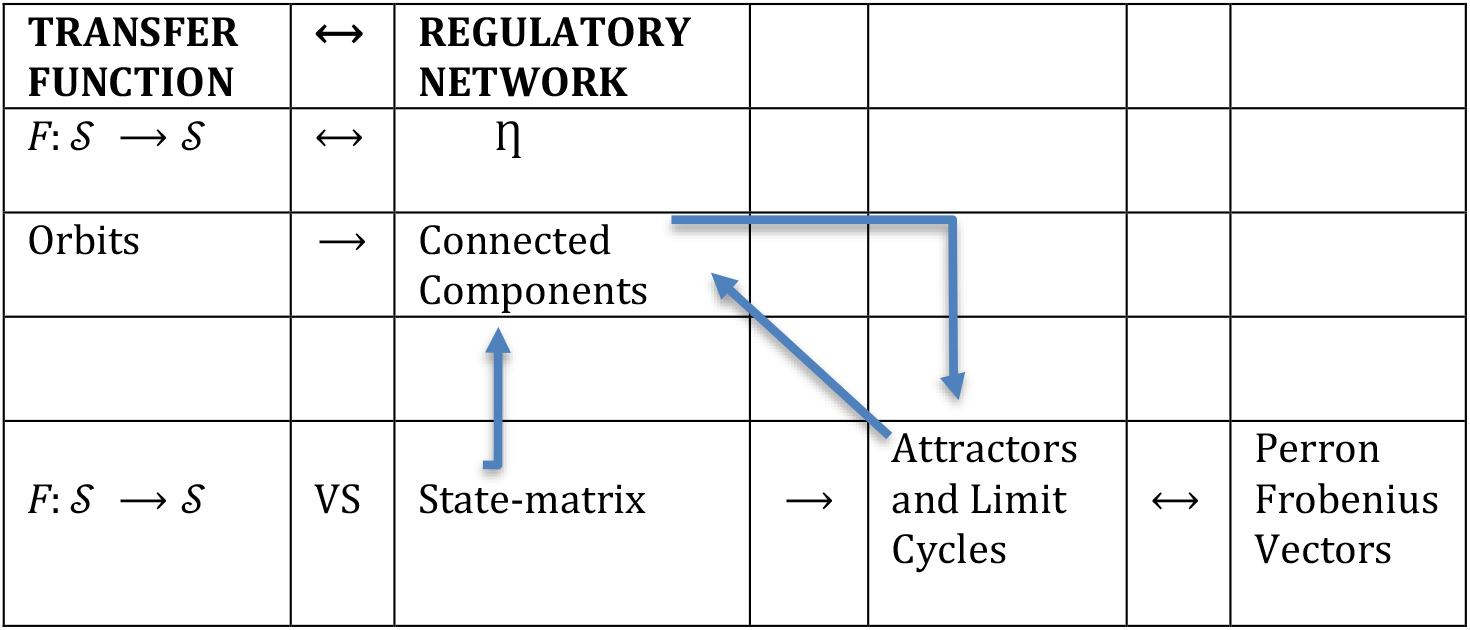
Information flow between *F* and η.

A more complex case shown in Fig 5b occurs when a dicycle in η contains more than one element connected in series and forming a loop. Such configuration forms a **limit cycle**, namely a closed loop that forms a stable feedback system. It must be noted that an attractor would be a limit cycle of size 1. Mathematically, there is a set of state-vectors, {*v*_1_, …, *v*_*m*_} with size *m*>1, such that:

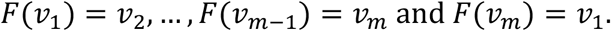

The limit cycle in the neural network of Tritonia swimming has been shown in Fig 2. It can be seen that it contains a peripheral element, a **ditree (T)**. Most regulatory networks in biology contain one or more ditrees, which form unidirectional connections to the main loop or dicycle of η. A connected component is an input that activates or modulates the dynamics of η. A network receiving ditrees is a **connected network**. Fig 5 exemplifies characteristic configurations of connected components with ditrees.

Having defined the set of components of the biological network, their connectivity and topology, the structure/function relationship can be quantitatively analyzed by understanding the information flow between F and η. Fig 6 summarizes the basis of such information flow.

Mathematically, a **connected component** of a network η is a maximal connected subgraph of its underlying graph |η|. For example, the network in Fig 4 has two connected components: *C*_*1*_ and *C*_2_. On the other hand, the network in Fig 2 is a connected network. For definitions and results of graph theory, see [18]). It must be noted that theoretically, η may be not connected, although most biological cases are connected networks with multiple inputs and regulatory elements.

### II.1. THE TRANSFER FUNCTION *F* AND ITS ORBITS

The regulatory network is built employing a set-function *F*: 𝒮 ⟶ 𝒮 that we will call the transfer function *F* and its compositions, as follows:

Let us recall that 𝒮 is the set of state vectors of the biological system of *n* elements (genes, cells, etc.). The time is important to the development of this work. Therefore, we can now confer a time-dependence to the evolution of the network as follows: *F*^0^= *I*_𝒮_ is the identity function of 𝒮, *F*^1^=*F*, and for each time *t* >1, *F* ^*t*^ =*F*∘*F*∘ ⋯ ∘ *F* denotes *t* times its composition. That is:

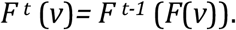

#### II.1.1

Since *F* is a set-function, see the glossary, for each *v* in 𝒮, there is a unique *w* such that, *v* is sent under *F* to *w*, as denoted by *F*(*v*)= (*w*). This property permits the connected components of the network to have some of the shapes described in Fig 6. From each element *v* in S emerges one unique arrow to *F(v)*. However, any element of S may receive two or more arrows.

#### II.1.2

The network η defined by *F* has as vertices the elements of 𝒮 and arrows *v*⟶ *F*(*v*), for each *v* in 𝒮.

The transfer function *F*: 𝒮 ⟶ 𝒮 is a set-function. Therefore, all its compositions *F*^*t*^ are also set-functions from 𝒮 to 𝒮. Since 𝒮 is a finite set with *s* elements, for each *v* in 𝒮, the set {*F*^*t*^(*v*): *t ≥*0} which is a subset of 𝒮 has at most *s* elements. Therefore, there exists 0≤*m < t* ≤ *s* such that *F*^*m*^(*v*)= *F*^*t*^(*v*) and thus *F*^*m*^(*v*)⟶ *F*^*m+1*^(*v*) ⟶ ⋯ ⟶*F*^*t*^(*v*)= *F*^*m*^(*v*) is a dicycle and by (II.1.1), this dicycle is unique.

#### II.1.3

We will now define an **orbit of a state-vector** *v*, whose interest relies on containing information about the evolution of the dynamics of the network. Mathematically, the orbit of *v* in 𝒮 is the set {*F* ^*t*^(*v*): *t ≥*0} and will be denoted by Ω(*v*). According to sections II.1.1 and II.1.2, Ω(*v*) contains a unique dicycle.

An example of topology with orbits and a ditree can be found in the regulatory network of Tritonia swimming (Fig 7), which has two different orbits, one of which incorporates all the elements:

**Figure 7.**
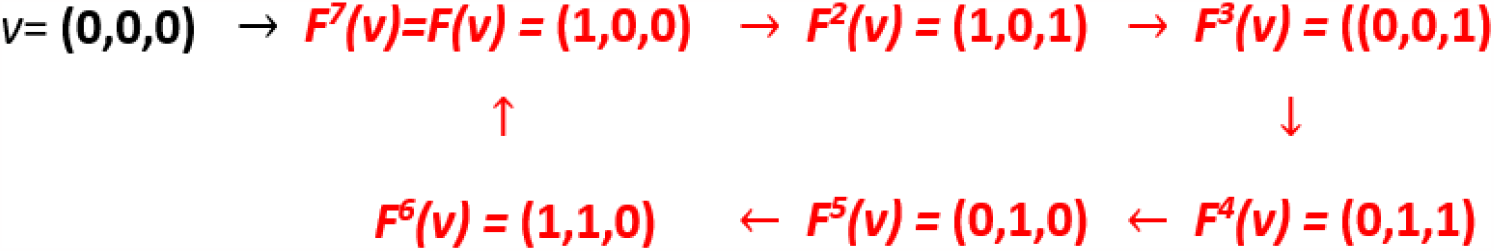
The network of Tritonia Swimming has an orbit that incorporates all the elements Ω(*v*) = {(0,0,0), (1,0,0), (1,0,1), (0,0,1), (0,1,1), (0,1,0), (1,1,0)} while the other orbits (marked in red) are the limit cycle of the network, (see the text).

**Figure 8.**
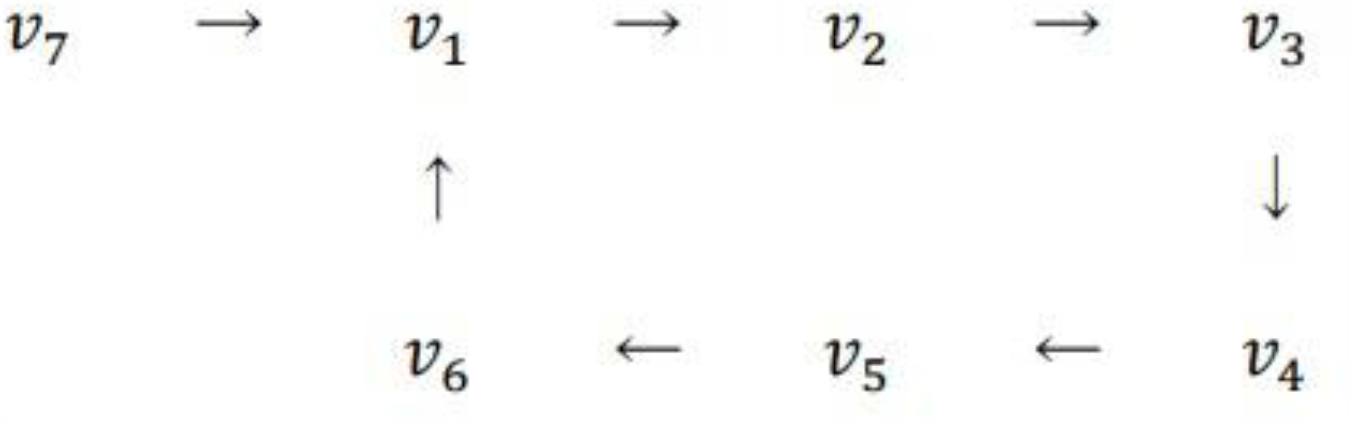
Structure of the regulatory network of Tritonia Swimming, with a unique ditree *v*_7_→*v*_1_.

Ω(*v*) = {(0,0,0), (1,0,0), (1,0,1), (0,0,1), (0,1,1), (0,1,0), (1,1,0)} with *v*=(0,0,0). The other orbit is the limit cycle of the network (red in Fig 7):

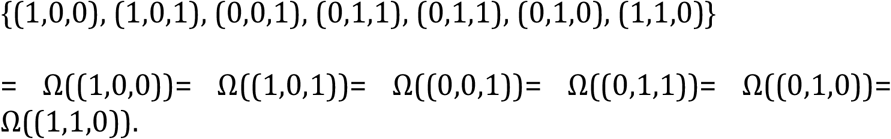

#### II.1.4

Since each orbit has a unique dicycle (as seen in II.1.3), any additional arrow is a ditree pointing to the dicycle. Fig 6 shows the general shape of the connected components of a regulatory network; sections II.4 and II.5 contain a method construct them. A computer routine for the same purpose is presented as a supplementary file.

### II.2. THE STATE-MATRIX OF η AND ITS SPECTRAL PROPERTIES

Now we will define the **state-matrix**, *M*_η_, which not only stores the information of the transition function see (II.1), but also gives us the temporal dynamics of the biological system.

To define *M*_η_ we must first fix an order in 𝒮 = {*v*_1_, …, *v*_*s*_}. *M*_η_ is a **real** matrix of size *s×s*, whose *ij*-th entry is:

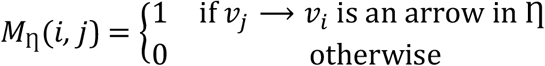

Any order renders equivalent biological information since the corresponding state matrices are conjugated.

Observe that 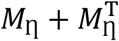 (where 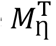 denotes the transpose matrix of *M*_η_) is the adjacency matrix of the underlying graph of η. As an example, we may now define the *M*_η_ of the Tritonia swimming regulatory network η (see II.1.3 or Fig2) as:

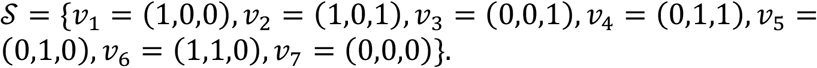

With such order, the state-matrix *M*_η_ of the network is:

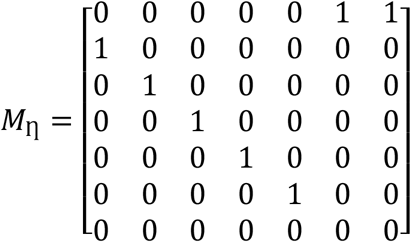

with eigenvector 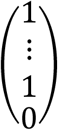 of the eigenvalue 1

#### II.2. Spectrum of *M*_η_

The characteristic polynomial of *M*_η_ gives the number of connected components of η. In addition, the Perron-Frobenius eigenvectors give the attractors and limit cycles. We will now see that the characteristic polynomials of the state matrices of the dicycles and ditrees fully describe the characteristic polynomial and the spectrum of the state matrix.

#### II.2.1

The characteristic polynomial *p*_σ_(*x*) of a dicycle *σ* of size *r*, is:

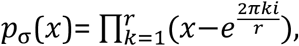

where 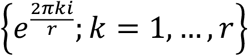 are the *r-*roots of unity, defined as the complex numbers *z*, such that *z*^*r*^=1, and *e*^2*πi*^ = 1 (see [18]).

The above result is known and it is also known that the state-matrix of a dicycle σ is as follows:

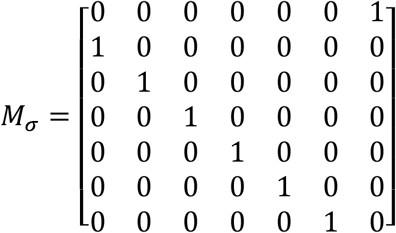

with eigenvector 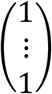 of the eigenvalue 1

#### II.2.2

The characteristic polynomial of a ditree *T* of size *q*, is *p*_*T*_(*x*)= *x*^*q*^ (see [18]). By convention (see II.1.3), a ditree *T* of η has a size bigger than 1 and contains a unique vertex in the dicycle of η (see Fig 7).

#### II.2.3

Now we may analyze the characteristic polynomial of *M*_η_ when the regulatory network η is connected. For definitions see the glossary. Recall that the vertices of η are the elements of 𝒮 = {*v*_1_, …, *v*_*s*_} and η has a unique dicycle σ of size *r* and possibly, *T*_1_, …, *T*_*d*_ ditrees with *d*≥0 and each *T*_h_ of size *q*_h_. Then the characteristic polynomial of *M*_η_ is:

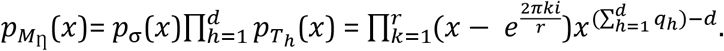

In this case, 1 is an eigenvalue of *M*_η_ with a unique (up to scalar multiples) eigenvector 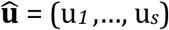, where u_*i*_ =1 if *v*_*i*_ belongs to the dicycle, and u_*i*_ =0 otherwise. Since all the eigenvalues have modulus equal to *r* ≤1 (and 1 is also an eigenvalue of *M*_η_), the maximum of the modulus of these eigenvalues is 1. This maximum is known as the **spectral radius** 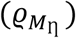 of *M*_η_ and is denoted by:

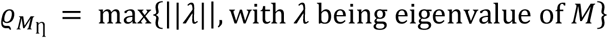

The eigenvector 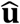 of 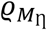 is the **Perron-Frobenius (eigen)vector** of *M*_η_ and gives us a way to find the vertices of η, which form its attractor or limit cycle, as will be shown in II.4.

#### II.2.4

If η is not connected, we can define an order of 𝒮 such that *M*_η_ has the following form:

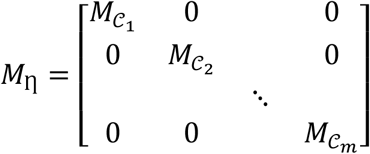

with 𝒞_1_, …, 𝒞_*m*_ being the different connected components of η (see the glossary) and their respective state-matrices being 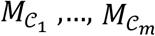(see II.0). Then, the characteristic polynomial of *M*_η_ is:

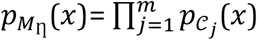

Since 𝒞_*j*_ is a connected network, 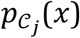 has the form described in II.2.3. Therefore, 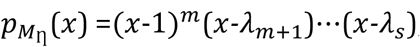, with *λ*_*m*+1_,*…, λ*_*s*_ ≠ 1. In other words, the maximum power of *x* − 1 in the characteristic polynomial of *M*_η_ gives the number of the connected components of η. Section II.5 contains an algorithm to find such components.

As an example, we may now take the network in Fig 4. Its characteristic polynomial is *x*^8^ − 2*x*^7^ + *x*^6^ = *x*^6^(*x* − 1)^2^ . The power of *x-*1 is 2, therefore as shown in Fig 4, η has two connected components.

### II.3. THE TRANSFER FUNCTION VERSUS THE STATE-MATRIX

To get more properties of the regulatory network, it will be useful to analyze the relationship between the transfer function *F* : 𝒮 ⟶ 𝒮 and the state-matrix *M*_η_, since together they store information about the network dynamics. While *F* informs about the individual state vectors *M*_η_ gives the “global connectivity” of each state vector.

Let *s* be the number of elements of 𝒮, and recall that for each *t*≥ 0, *F*^0^ = *I*_𝒮_, *F*^1^ = *F* and *F*^*t*^ = *F* ∘^…^∘ *F t* times the composition of *F*, and *M*_η_ is a real matrix of size *s×s* (See II.1).

For each subindex *i* = 1, …, *s*, denoted by *e*_*i*_ the real column-vector having 1 in the *i*-th coordinate and 0 elsewhere. Now we have two types of bijective set functions The first type occurs between the set of state-vectors 𝒮 and the assembly of vectors {*e*_1_, …, *e*_*s*_}:

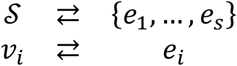

Recall that any real vector is a linear combination of the vectors *e*_1_, …, *e*_*s*_. Therefore, to study the state-matrix *M*_η_ it is enough to study the images of these vectors under *M*_η_ . The second type of bijection, is established between the orbit of *v*_*i*_ under *F*, {*F*^*t*^*v*_*i*_: *t* ≥ 0}, and the orbit of *e*_*i*_ under the matrix *M*_η_, 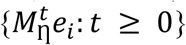. Recall that any real vector is a linear combination of the vectors *e*_1_, …, *e*_*s*_. Therefore, for each state-vector *v*_*i*_, we have one bijection:

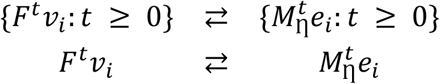

### II.4. THE CONNECTED COMPONENTS OF THE NETWORK η

We can now discuss a method to find the connected components of η. As we mentioned before, each connected component must be studied separately to avoid mixing information that will render false results.

#### II.4.1

As described in II.2.4, the characteristic polynomial of *M*_η_ is 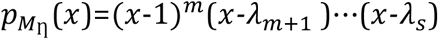, where *λ*_*m*+1_,*…, λ*_*s*_ are different from 1. Therefore, η has exactly *m* different connected components. Moreover, the nonzero entries of the eigenvectors associated with the eigenvalue 1 are the vertices of the attractors and limits cycles of η.

Following the example in Fig 4 and (II.2.4), the vertices of η are:

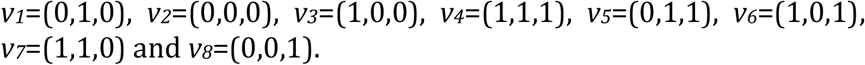

The characteristic polynomial of *M*_η_ is 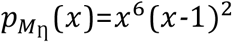, with the power of *x-*1 being 2. Therefore, η has two connected components. The eigenvectors of 1 that give the vertices in attractors and limit cycles are (0,1,0,0,0,0,0,1)^*T*^ and (0,0,0,0,0,0,0,1)^*T*^, with their non-zero entries corresponding to the state-vectors *v*_*2*_ and *v*_*8*_. Now to get the different attractors and limit cycles we calculate their orbits Ω(*v*_*2*_) = {*v*_*2*_} and Ω(*v*_*8*_) = {*v*_*8*_}. Therefore, η has exactly two attractors, one in *v*_*2*_ =(0,0,0) and the other in *v*_*8*_ =(0,0,1), as we can confirm in the Fig 4.

#### II.4.2

Having the attractors and limit cycles of the network η, we need to find the vertices in the connected components they define.

Using orbits again, if a vertex *v* is in a connected component *C*, the complete orbit Ω(*v*) is contained in *C* and Ω(*v*) contains the dicycle of *C*. Therefore, two state vectors belong to the same connected component if their orbits have a non-empty intersection. Moreover, they must share the same dicycle.

Following the example in Fig 4 and II.4.1, we get that the vertices *v*_*1*_=(0,1,0), *v*_*3*_=(1,0,0), *v*_*4*_=(1,1,1), *v*_*5*_=(0,1,1), *v*_*6*_=(1,0,1), *v*_*7*_=(1,1,0), and *v*_*8*_=(0,0,1) form a connected component, while the other connected component is a loop in *v*_*2*_=(0,0,0), as one can verify in II.2.4.

### II.5. AN ALGORITHM TO CONSTRUCT THE CONNECTED COMPONENTS OF η

We are now in a position to reconstruct the dynamics of the network from the following cookbook, which is algorithmizable in every step:

1. Obtain *M*_η_ using any order of 𝒮.
2. η has *m* different connected components, where *m* is the maximal power of *x*-1 in the characteristic polynomial 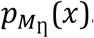. That is: 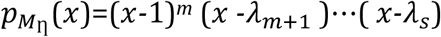, with *λ*_*m*+1_,*…, λ*_*s*_ different from 1. The set of vertices corresponding to the non-zero coordinates of the eigenvectors of eigenvalue 1 are the vertices of the dicycles of η. Now take one of those vertices, say *v*, then the orbit Ω(*v*) must be the attractor or limit cycle of a connected component of the network η.
3. Now we will build each connected component. The *m* different dicycles say σ_1_, …, σ_*m*_, correspond to *m* different connected components, we have to find their state vectors. By (II.4.2), two state vectors belong to the same connected component if their orbits have a non-empty intersection.

### III. TOPOLOGY OF THE NETWORK

This section provides mathematical evidence as to why the attractors and limit cycles inform about the temporal dynamics of the biological system. The technique will be illustrated in the Tritonia regulatory network, Figs 1, 2 and 7.

#### III.1

The regulatory network of the Tritonia swimming η is connected (see the glossary) and consists of seven state vectors, see Fig 7, where the state vectors of its limit cycle are:

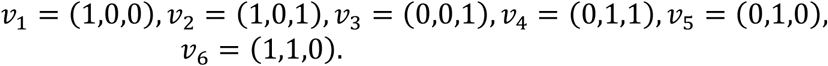

The 7 by 7 state matrix and its eigenvector û with eigenvalue 1 have been shown in section II.1. Note that η has a unique ditree *v*_7_→*v*_1_ with *v*_7_ = (0,0,0). The *ij-*th entry of *M*_η_ occurs into the *i*th row and the *j*th column, and has a 1 value if there is an arrow in η from *v*_*j*_ to *v*_*i*_.

As described in sections II.2 and II.3, the *i*th entry corresponds to the state-vector *v*_*i*_, which has a value of 1, corresponding to a state-vector in a limit cycle. Moreover, 1 is the spectral radius of *M*_η_ and its eigenvector is the Perron-Frobenius vector of the state-matrix.

As we will see in (III.5), for some matrices, for example for *M*_η_ when η is a connected network (see the glossary for definitions), the Perron-Frobenius vector has the following asymptotic property:

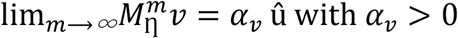

For every *v* in 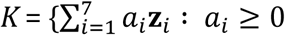 for *i*=1,…,7}. This set *K* is built with the orbits of the state-vectors of the network, see (II.1.3) and (II.3). In this case of the Tritonia, the vectors **z**_*i*_ are the following:

**z**_**1**_=(0,1,1,1,1,1,0) vector associated to the orbit *v*_1_.

**z**_**2**_=(1,0,1,1,1,1,0) vector associated to the orbit *v*_2_.

**z**_**3**_=(1,1,0,1,1,1,0) vector associated to the orbit *v*_3_.

**z**_**4**_=(1,1,1,0,1,1,0) vector associated to the orbit *v*_4_.

**z**_**5**_=(1,1,1,1,0,1,0) vector associated to the orbit *v*_5_.

**z**_**6**_=(1,1,1,1,1,0,0) vector associated to the orbit *v*_6_.

**z**_**7**_=(1,0,0,0,0,0,1) vector associated to the orbit *v*_7_.

Fig 9 explains the meaning of the limit just described.

**Figure 9.**
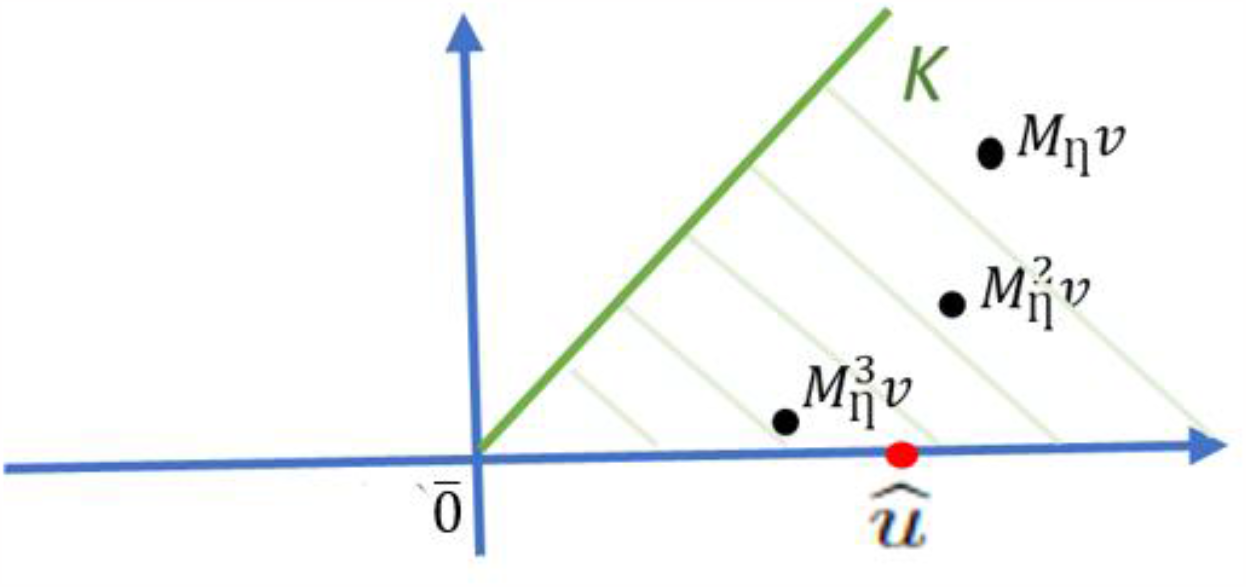
Geometric interpretation of the Birkhoff-Vandergraft theorem described in section III.5. 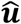 is the Perron-Flobenius vector; *K* is the set of real vectors *v*, and 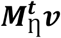 is the state matrix of η for a vector *v* over time *t*.

Now we will put all the previous facts into context.

#### III.3

According to the Jordan decomposition theorem, the eigenvalues and eigenvectors of a matrix control the behavior of the linear function defined by the matrix (see [22,23]). More specifically, the spectral radius of the state-matrix *M*_η_ provides an asymptotic prediction of the behavior of the connected network over time (see sections III.1, III.2 and III.5).

The set *K* of real vectors is defined by the orbits of the state vectors of η (see II.1.3). *K* is invariant under the state-matrix *M*_η_, meaning that for every vector *v* in *K, M*_η_*v* also lies in *K*. Moreover, for *t* ≥ 0, 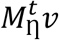 approaches the ray defined by the Perron-Flobenius eigenvector vector 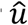, with its unique eigenvalue *ρ*(*M*_η_) = 1 (see II.2.3). Such ray is the asymptote of 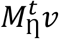 which informs about the temporal dynamics of the biological network.

#### III.4

To construct the set *K*, we must first recall that the vertices of η are the state vectors 𝒮 = {*v*_1_, …, *v*_*s*_} with {*v*_1_, …, *v*_*m*_} being the dicycle or the network. If *m*=1, {*v*_1_, …, *v*_*m*_} represents the attractor. Otherwise, {*v*_1_, …, *v*_*m*_} represents the limit cycle of η.

Now we must define the following real vectors in *s* coordinates:

For 1≤ *i* ≤ *s*, the vector **z**_***i***_ defined below, contains the information of the state-vectors in the orbit of the state-vector *v*_*i*_, as follows:

Its *j*th*-*coordinate is:

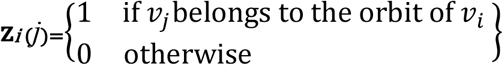

Recall that the orbit of *v*_*i*_ is:

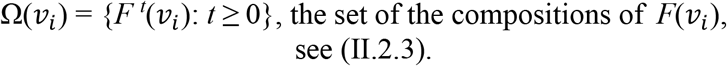

We can now define *K* as the set of nonnegative linear combinations of **z**_**1**_, …, **z**_***s***_.,

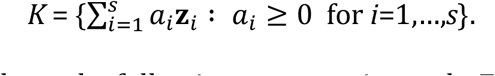

#### III.5

We now have the following asymptotic result: For every *v* in *K*, the following limit exists:

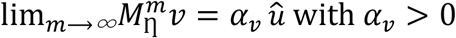

The above result emerges from the Birkhoff-Vandergraft Theorem (see [24]), which is a generalization of the Perron-Frobenius theorem (see [22,23]).

Therefore, 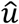 predicts the future behavior of the biological system. In other words, 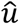 is truly a “limit vector”. Moreover, by (II.2), the non-zero entries of 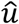 correspond to the state-vectors of the limit cycle or attractor of the network.

### III.6. ALGORITHM TO FIND THE ATTRACTORS AND THE LIMIT CYCLES OF η

If a biological regulatory system is small, it is not difficult to find its attractors and limit cycles. However, if we don’t know if the network is connected, finding its attractors or limit cycles is not an easy task. The following is an algorithm to find them:

1. Find all the connected components of η (see II.5).
2. Let 𝒞_1_, …, 𝒞_*m*_ be the different connected components of η and 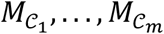 their respective state-matrices.
3. For each *h=*1, …, *m*, calculate an eigenvector of 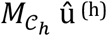 with eigenvalue 1. The state-vectors of 𝒮 associated with the nonzero coordinates of û ^(h)^ form the limit cycle of 𝒞_h_.

## Conclusions

1. We present a method to identify and represent a diversity of interactions among the different components that determine the working of regulated biological networks.
2. A logical digraph of the regulatory biological network is built by use of eight logical connectives designed from Boolean algebra, which in combination represent accurately every possible interaction among elements of the network.
3. Rules are provided to translate from logical to Boolean representation of the components of the network.
4. The transfer function, its orbits and the state matrix of the network allow us to determine the topology of the network, its limit cycles, attractors and dynamics.
5. The spectrum of the state matrix renders the Perron-Flobenius eigenvalues, which predict the time evolution of the network.
6. An algorithm and three software routines are provided to find the attractors and limit cycles of a biological regulatory system, to build its state matrix and to find the eigenvectors which are difficult to find in systems with a large number of elements.

## Acknowledgment

We wish to express our gratitude to Felipe Meneses for his help in obtaining valuable bibliographical material, to Isabel Takane for the typography of the article, and to Francisco Pérez Eugenio for continuously supporting our computer requirements.

## SUPPORTING INFORMATION

### COMPUTATIONAL PROGRAMS

These programs permit to obtain the regulatory network of a biological system and its connected components. Recall that in a biological system with *n* elements: 𝒢 ={*g*_*1*_,*…,g*_*n*_} is an **ordered** set of genes, neurons, cells or nodes. 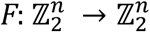 is the transition function. *F=*(*f*_*1*_,*…,f*_*n*_) is given by *n* Boolean functions 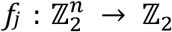. η is the regulatory network whose vertices are the elements of the set of state-vectors 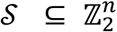 and for each state-vector an arrow (*a*_1_, …, *a*_*n*_) → *F*(*a*_1_, …, *a*_*n*_).

#### S1. WOLFRAM ALGORITHM TO BUILD FROM *F*, THE CONNECTED COMPONENTS OF η

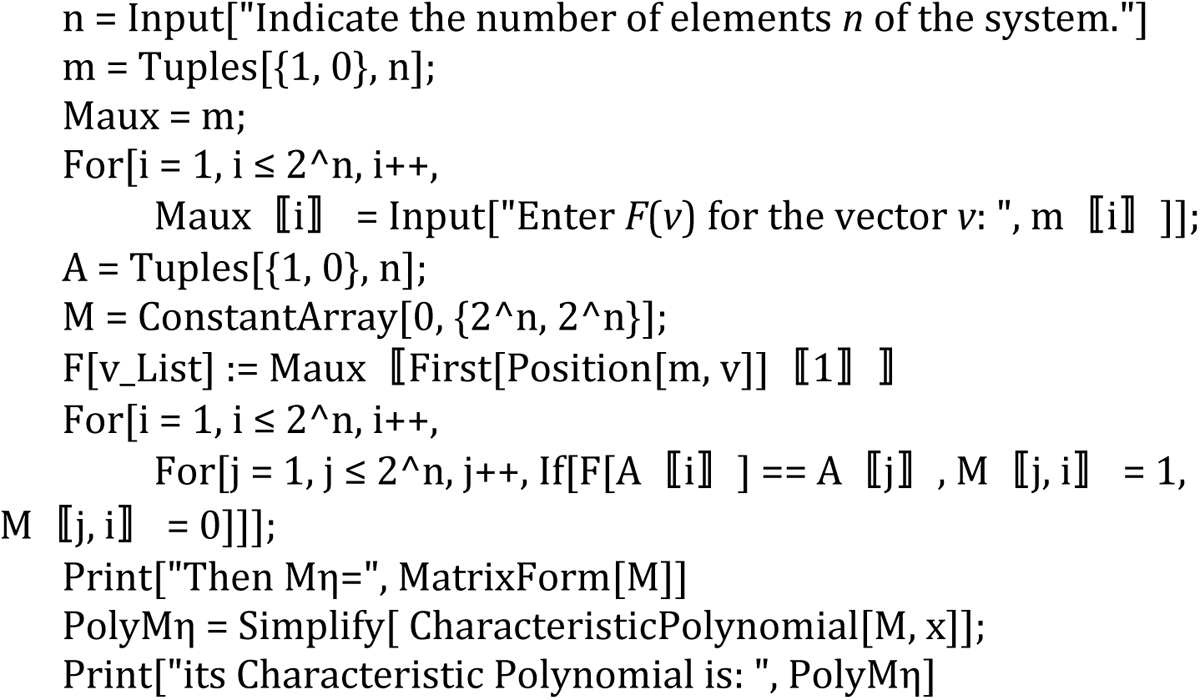
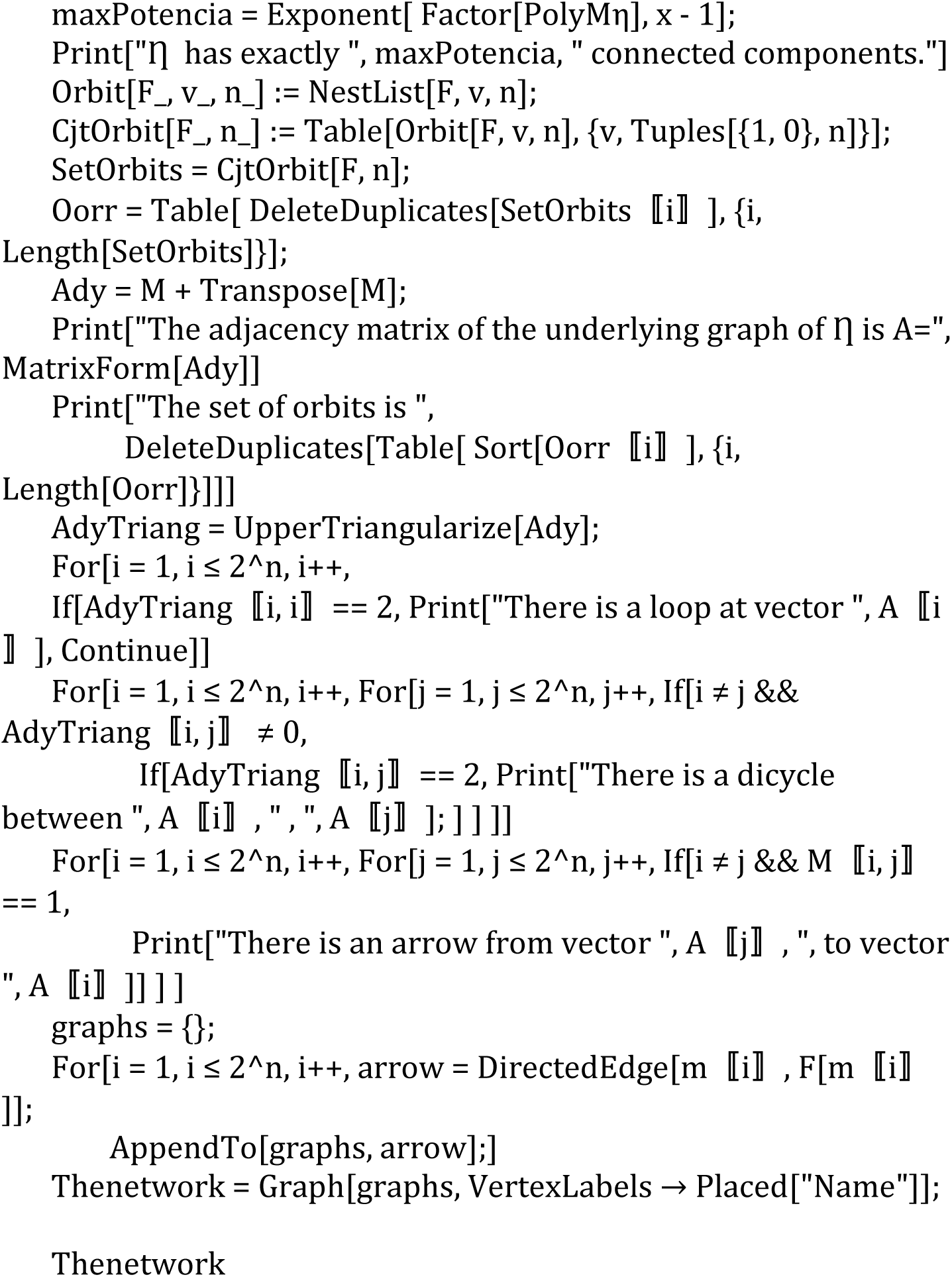

**S2**. Alternatively, from η and using the first part of this paper, we can obtain the transfer function *F* of the biological system. Here is a computational program to obtain *F*.

#### SM2. WOLFRAM ALGORITHM TO BUILD THE TRANSITION FUNCTION *F* FROM THE NETWORK η

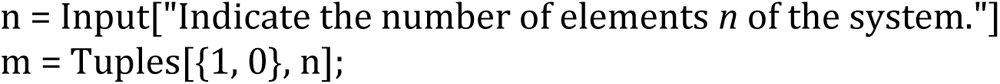
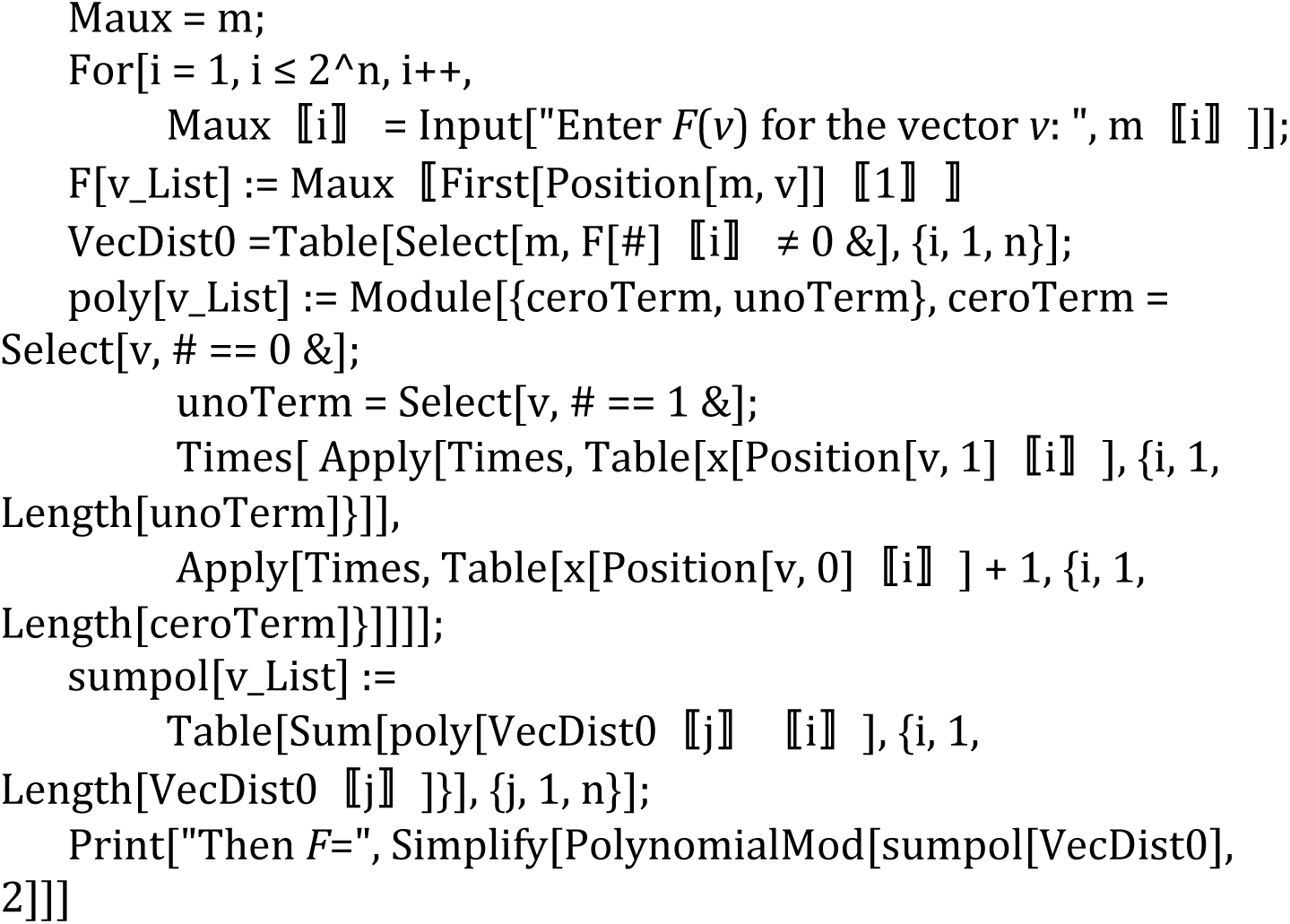

